# FLASH radiotherapy using high-energy X-rays: validation of the FLASH effect triggered by a compact single high-energy X-ray source device

**DOI:** 10.1101/2024.07.16.603758

**Authors:** Binwei Lin, Huan Du, Yiwei Yang, Xiaofei Hao, Feng Gao, Yuwen Liang, Wenqiang Tang, Haonan Xu, Mingming Tang, Yao Liao, Decai Wang, Bo Lin, Yihan Zhu, Tingting Wang, Runqiu Gu, Xin Miao, Yixiao He, Yu Zhang, Jie Li, Zheng Zhou, Jianxin Wang, Dai Wu, Xiaobo Du

**Affiliations:** Mianyang Central Hospital, School of Medicine, University of Electronic Science and Technology of China, Mianyang, 621000, China; NHC Key Laboratory of Nuclear Technology Medical Transformation, Mianyang Central Hospital, School of Medicine, University of Electronic Science and Technology of China, Mianyang, 621000, China; Sichuan Clinical Research Center for Radiation and Therapy, Mianyang Central Hospital, Mianyang, 621000, China; Institute of Applied Electronics, China Academy of Engineering Physics, Mianyang, 621900, China; Clinical Medical School, North Sichuan Medical College, Nanchong 637000, P.R. China; National Key Laboratory of Science and Technology on Advanced Laser and High Energy Microwave Mianyang, 621000, China

**Keywords:** FLASH radiotherapy, FLASH effect, CHEXs, Single gantry rotation, short-interval fractional

## Abstract

**Purpose:** This preclinical study aimed to verify the FLASH effect of compact single high-energy X-ray source (CHEXs) and to explore whether three irradiations with single-gantry rotation two 30 s pauses can generate FLASH effect in mice.

**Materials and methods:** The absolute dose and pulsed beam of the CHEXs were measured using an EBTXD radiochromic film and fast current transformer. Healthy C57BL/6J female mice and a subcutaneous tumor model were irradiated under different conditions: sham (control), FLASH-RT (FLASH1: delivering the total dose in 1 fraction; FLASH3: delivering the total dose with two 30 second pauses to simulate a three-field delivery where the gantry rotation is occurring within 30 seconds), and conventional dose rate radiotherapy (CONV-RT). Various total doses were administered to the corresponding normal tissues (whole thorax, 30 Gy; whole abdomen, 12 Gy; and skin, 36 Gy) and tumors (CT26, 16.5 Gy; and LLC, 18 Gy). Survival status, normal tissue damage, and tumor growth suppression were recorded in each group.

**Results:** The average dose rate of the CHEXs exceeded 40 Gy/s. For whole-thorax and skin irradiation, both FLASH1 and FLASH3 demonstrated protective effects. For whole-abdomen irradiation, FLASH1 exhibited a superior protective effect. No significant differences in tumor growth responses were observed between the FLASH1, FLASH3, and CONV-RT groups (P>0.05).

**Conclusion:** This study confirmed that the FLASH effect could be triggered using CHEXs FLASH radiotherapy, and demonstrated that three irradiations with single gantry rotation two 30 s pauses can trigger the FLASH effect, indicating the potential benefit of CHEXs 3D conformal radiotherapy. Our findings indicate that further clinical trials on CHEXs are warranted.

## Introduction

Cancer remains a leading cause of death worldwide, with an estimated 20 million new cases and approximately 9.7 million deaths reported in 2022 [1]. Radiotherapy, a commonly used treatment for cancer, is required by approximately 60–80% of affected patients throughout the course of the disease [2,3]. However, radiotherapy can damage the surrounding organs at risk (OARs), resulting in treatment interruption and impaired anti-tumor efficacy. Moreover, radiotherapy-related toxicities, such as radiation pneumonia, enteritis, and skin damage, can lead to respiratory failure, diarrhea, and infection, significantly affecting the patient’s quality of life and survival [4]. Therefore, exploring methods to reduce these toxicities is a critical focus of research.

Ultra-high dose rate radiotherapy (FLASH-RT) refers to a radiotherapy technique that uses dose rates exceeding 40 Gy/s to treat cancer [5], compared with conventional dose rate radiotherapy (CONV-RT), which use dose rates ≤0.1 Gy/s. The primary advantage of FLASH-RT is that it can minimize OARs damage while maintaining anti-tumor efficacy [5, 6], a phenomenon known as the FLASH effect. Moreover, its higher OAR tolerance allows transmission of higher FLASH-RT doses to the tumor site, potentially further improving treatment efficacy for radiation-resistant tumors. In addition, FLASH-RT can be completed within a very short time[7], reducing patient time consumption and economic burden. Electrons [6,8], protons [9,10], and photons [11,12] are the most commonly used radiation sources in FLASH-RT. Generally, electron beams are only suitable for the treatment of superficial tumors, such as skin tumors, given their limited penetration depth. Although proton beams may be suitable for deep tumors, their high construction and operating costs limit their application. Alternatively, high-energy X-rays, with deep penetration, minimal divergence, and affordability, are the most promising candidates for use in FLASH-RT.

Our previous study established the PARTER platform, a preclinical research platform, on the China Academy of Engineering Physics THz free electron laser (CTFEL), capable of generating high-energy X-rays at ultrahigh dose rates (700–1200 Gy/s, 6–8 MV) that we found were capable of triggering a FLASH effect by effectively mitigating acute radiation damage to lung and intestinal tissues without compromising anti-tumor effects [12]. However, the CTFEL is a preclinical experimental platform that is bulky and expensive, making it unsuitable for further clinical research due to its size. Therefore, to develop a high-energy X-ray FLASH radiotherapy equipment suitable for clinical use, we developed a compact single high-energy X-ray source (CHEXs) FLASH radiotherapy device based on ambient-temperature linear accelerator technology. CHEXs can be directly installed within existing medical linear accelerator rooms without expansion, with a diameter similar to conventional equipment (approximately 3.1 m). Moreover, the CHEXs device uses an alternative pulse mode (S-band), which combines economic efficiency and small size to accelerate electrons, achieving high-energy electron beams (≤ 10MeV). These beams bombards high-speed rotating tungsten targets, converting them into ultra-high dose rate X-rays via bremsstrahlung radiation [13].

Although our previous study verified that high-energy X-rays at ultra-high dose rates can induce the FLASH effect, other studies have suggested that dose rate alone is not the sole determinant, with additional physical parameters, such as pulse structure [18–21] and single dose [22–24], potentially also influencing FLASH effect occurrence. Therefore, since the S-band used in CHEXs differs from that of the PARTER system, this preclinical study aimed to verify the FLASH effect of CHEXs and to explore whether three irradiations with single-gantry rotation two 30 s pauses irradiation can generate FLASH effect, thereby potentially replacing multi-accelerator synchronous irradiation and providing a theoretical basis for subsequent clinical trials.

## Materials and Methods

### Irradiation device and dosimetry

All FLASH-RT experiments were conducted using the CHEXs device (Mianyang, China), which can achieve an average dose rate of 81.01 Gy/s at a source-to-surface distance (SSD) of 1 m [13]. All CONV-RT experiments were performed using a clinically applied 6 MV Elekta Precise linac (Elekta AB, Stockholm, Sweden). Dose monitoring procedures were consistent with those described previously [12]. Beam current was monitored using a bushing-current transformer (BCT), with a diamond detector mounted downstream of the primary collimator for X-ray beam monitoring. Briefly, GafchromicTM EBT-XD radiochromic films (Ashland Inc., Covington, Kentucky, USA) were placed beneath solid water at the central level of the irradiation target area to ensure uniform absolute dose irradiation. Before the FLASH experiment, the EBT-XD film was calibrated using a clinically applied 6 MV Elekta Precise linac. Figure 1A presents the schematic of the in vivo FLASH-RT experiment. Lead secondary collimators with apertures of 8 × 4 cm^2^, 4 × 4 cm^2^, 3 × 2 cm^2^, and 2 × 1.2 cm^2^ were used to delimit the FLASH irradiation field (Fig. 1B). The total irradiation dose was controlled by varying the exposure time. Figures 1C and 1D illustrate the fixation and whole-thorax irradiation setup for mice in the FLASH-RT and CONV-RT groups. During FLASH-RT, EBT-XD films were placed on the anterior surface of each mouse to monitor the total dose of a single irradiation event (Fig. 1C). For whole-thorax irradiation of CONV-RT, a 2 cm wide strip field was used, and a 1.5 cm water-equivalent material was applied to the front of the mouse for dose build-up (Fig. 1D). EBT-XD films were placed at depths of 1.5 cm and 2.5 cm under the solid water to simulate the dose at the center of the whole-thorax irradiation target to ensure consistent irradiation dose for both the FLASH-RT (Fig. 1E and 1G) and CONV-RT (Fig. 1F and 1G) groups. The percent depth dose (PDD) curves comparing the FLASH-RT and CONV-RT groups are shown in Figures 1H, 1I, and 1J. Additional information on whole-abdomen, skin and tumor irradiation can be found in Supplementary Fig. 1.

**Fig. 1:**
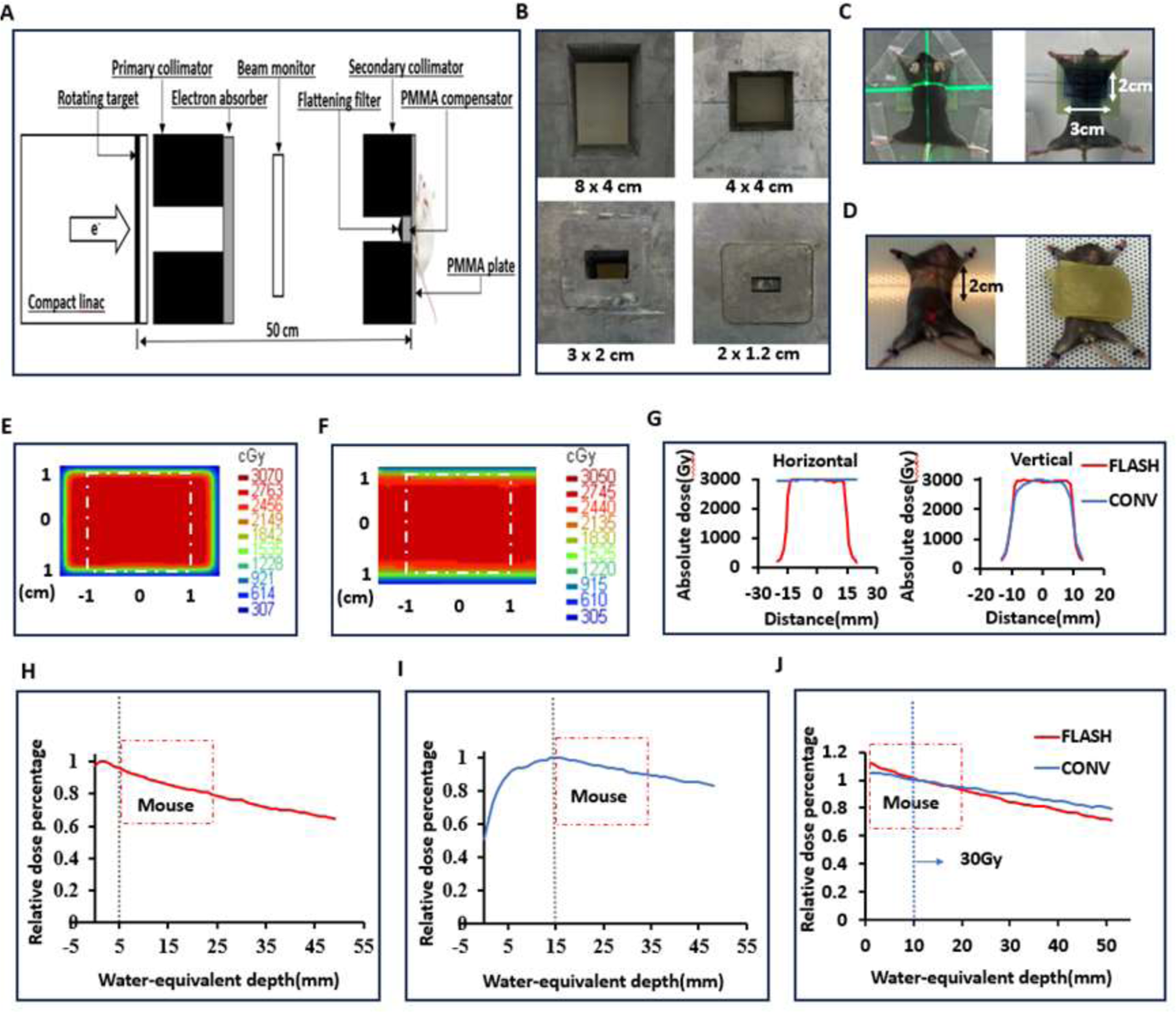
Parameters and dosimetry of FLASH-RT and CONV-RT. A: Schematic diagram of the in-vivo FLASH X-ray experiment. B: Secondary collimators (aperture, 8 × 4 cm^2^, 4 × 4 cm^2^, 3 × 2 cm^2^, 2 × 1.2 cm^2^) used for delimitating the FLASH irradiation field. C: For whole-thorax FLASH-RT, a 5-mm thick polymethyl methacrylate (PMMA) plate was used for dose build-up and mice fixing, with an EBTXD film placed between the PMMA plate and the anterior surface of the irradiated mouse for dose evaluation. D: A 2-cm wide strip field used for whole-thorax CONV-RT, with a 1.5 cm water equivalent material covering the front of the mouse for dose build-up. E: Dose distribution of an EBTXD film mounted at 1.5-cm depth in solid water (calibration dose at the central level of FLASH-RT mice). F: Dose distribution of an EBTXD film mounted at 2.5-cm depth in solid water (calibration dose at the central level of CONV-RT mice). G: The horizontal and vertical dose profiles, representing the dose distribution in FLASH-RT (E) and CONV-RT (F). H: The PDD curve of X-ray sources used for FLASH-RT, with the mouse (red dashed box) placed 5 mm from the secondary collimator with a 5-mm build-up area depth. I: The PDD curve of X-ray sources used for CONV-RT, with the mouse (red dashed box) placed 15 mm from the secondary collimator with a 15-mm built-up area depth. J: A 30 Gy administered 1 cm under the skin, with comparable PDD curves between the FLASH-RT and CONV-RT groups.

### Mice and ethics statement

Female C57BL/6J and BALB/c mice, aged 6–8 weeks, were procured from Sibeifu Experimental Animal Technology Co., Ltd. (Beijing, China). Mice were subjected to irradiation exposure at 7 to 9 week of age. All animal experiments were conducted in accordance with the relevant ethical guidelines and approved by the Animal Ethics Committee of Mianyang Central Hospital (approval number: P2020032).

### Protective effects on normal tissues

For whole-thorax irradiation, the C57BL/6J mice were randomly divided into four groups: control (0 Gy; n=18), FLASH1 (30 Gy × 1 F, 10 MeV; n=18), FLASH3 (10 Gy ×3 F, 30 s interval, 10 MeV; n=15), and CONV (30 Gy ×1 F, 6 MeV; n=18). The entire thorax of each mouse was irradiated using a field size of 2 cm (craniocaudal) × 3 cm (lateral), with the upper boundary of the irradiation field located below the edge of the ear. The dose rates for FLASH-RT (FLASH1 and FLASH3) and CONV-RT were 340 Gy/s and 0.07 Gy/s, respectively. At 72 h post-irradiation, five mice from each group were euthanized, and lung tissues fixed in 4% paraformaldehyde were collected for hematoxylin-eosin (H&E) staining. The survival status of the remaining mice was monitored. The endpoints for survival included death, weight loss>20% and self-harm.

For whole-abdomen irradiation, the C57BL/6J mice were randomly divided into four groups: control (0 Gy; n=15), FLASH1 (12 Gy ×1 F, 10 MeV; n=15), FLASH3 (4 Gy × 3 F, 30 s interval, 10 MeV; n=15), and CONV (12 Gy × 1F, 6 MeV; n=15). The entire abdomen of each mouse was irradiated using a field size of 4 cm (craniocaudal) × 4 cm (lateral), with the upper boundary of the irradiation field located at the lower edges of the lungs (2 cm below the bilateral ear edges). The dose rates for FLASH-RT (FLASH1 and FLASH3) and CONV-RT were 244 Gy/s and 0.07 Gy/s, respectively. At 72 h post-irradiation, five mice in each group were euthanized, and small intestine tissues (the jejunum at a distance of 5cm from the stomach, with a total length of 5cm) fixed in 4% paraformaldehyde was collected for H&E staining. The survival status of the remaining mice was determined. The endpoints for survival included death, weight loss>20% and self-harm.

For skin irradiation, the C57BL/6J mice were randomly divided into four groups: control (0 Gy; n=8), FLASH1 (36 Gy × 1 F, 10 MeV; n=8), FLASH3 (12 Gy × 3 F, 30 s interval, 10 MeV; n=7), and CONV (36 Gy × 1F, 6 MeV; n=7). The thigh skin of each mouse was irradiated using a field size of 2 cm (craniocaudal) × 3 cm (lateral), with the upper boundary of the irradiation field at the inguinal mouse region. The dose rates for FLASH-RT (FLASH1 and FLASH3) and CONV-RT were 350 Gy/s and 0.07 Gy/s, respectively. Photographs of the irradiated area on each mouse were taken every Monday and Thursday post-irradiation. Radiation dermatitis severity was independently assessed using a four-grade scale based on a previously published study [14] by two trained animal researchers. Briefly, grade 0: nomal mouse skin; grade 1: dry desquamation; grade 2/3: moist desquamation/ulceration. Any inconsistent grading was discussed before determining the final grading. At 8 weeks post-irradiation, all mice were euthanized, and skin samples from the irradiated area were fixed in 4% paraformaldehyde for H&E and Masson’s trichrome staining.

### Tissue staining and evaluation

The H&E and Masson’s trichrome staining protocol were performed according to a previously published study [15]. H&E staining under light microscopy was used to evaluate radiation-induced damage in the lung, intestine, and skin tissues. For intestinal slices, the number of regenerating crypts per single field of view (40x magnification) and height of small intestinal villi were quantified. For skin slices, ImageJ software (National Institutes of Health, USA) was used to evaluate the epidermal thickness and degree of fibrosis, defined as the percentage of blue-stained area within a single field of view (10x magnification) on a Masson’s trichrome-stained section [11].

### Anti-tumor effects

Mouse lung cancer cells (LLC) and mouse colon cancer cells (CT26) were purchased from ATCC (Manassas, Virginia, USA). Cell cultures were maintained under standard conditions (37°C, 5% CO_2_), and tumor inoculation was performed after obtaining sufficient cell numbers [16]. For LLC cells, C57BL6J mice were inoculated in the right hind flank with 5 × 10^5^ cells. For CT26 cells, BAL b/c mice were inoculated in the right hind flank with 5 × 10^5^ cells. Tumors were irradiated when the tumor volume reached 30–100 mm^3^.

Subcutaneous tumor models (LLC, and CT26 cells) were randomly divided into four groups: control (LLC: 0 Gy, CT26: 0 Gy; 8-9 mice per group), FLASH1 (LLC: 18 Gy x 1 F, CT26: 16.5 Gy × 1 F; 10 MV; 8-9 mice per group), FLASH3 (LLC, 6 Gy × 3 F, CT26: 5.5 Gy × 3 F; 10 MV; 8-9 mice per group), and CONV (LLC: 18 Gy × 1 F, CT26: 16.5 Gy × 1 F; 6 MV; 8-9 mice per group). A field size of 1.2 cm (craniocaudal) × 2 cm (lateral) was used to irradiate the tumors. The dose rates for FLASH-RT (FLASH1 and FLASH3) and CONV-RT were 244 Gy/s and 0.07 Gy/s, respectively. Tumor volume was measured twice per week by the same researcher using a digital caliper and calculated using the following formula for an oblate ellipsoid: (length × width^2^)/2 [17]. Mice were euthanized when the tumor volume reached ≥2000 mm^3^ or if weight loss exceeded 20%. Tumor tissues were fixed in 10% formalin and evaluated for tumor necrosis using H&E staining.

### Statistical analysis

All statistical analyses were performed using SPSS 22.0 (IBM Corp., Armonk, New York, USA) or GraphPad Prism 8.0 (GraphPad Software, Inc., La Jolla, California, USA). The results are expressed as mean ± standard deviation or as proportions. Group comparisons were performed using one-way analysis of variance. Survival analysis was conducted using the Kaplan–Meier method, and differences between groups were assessed using the log-rank test. P-values <0.05 were considered statistically significant and all tests were two-sided.

## Results

The experimental parameters for whole-thorax, whole-abdomen, skin, whole-body, and tumor irradiation are listed in Supplementary Table 1. The average dose rate in this study ranged from 244 to 350 Gy/s.

At 98 days post-whole-thorax irradiation, the survival rates were 100%, 86.4%, 50%, and 0% in the control, FLASH1, FLASH3, and CONV groups, respectively (Fig. 2A). Survival rates were significantly lower in the CONV group than those in the FLASH1 (P<0.001) and FLASH3 (P=0.001) groups, and lower in the FLASH3 group than that in the FLASH1 group (P=0.081). In the lung tissues, H&E sections at 72 h post-irradiation in the CONV group revealed a significant decrease in the number of alveoli, widened alveolar septum, lung tissue hemorrhage, and inflammatory cell infiltration (Fig. 2B). The lung tissue structure in the FLASH1 group was similar to that of the control group, while the degree of damage in the FLASH3 group was between those of the CONV and FLASH1 groups (Fig. 2B).

**Fig. 2:**
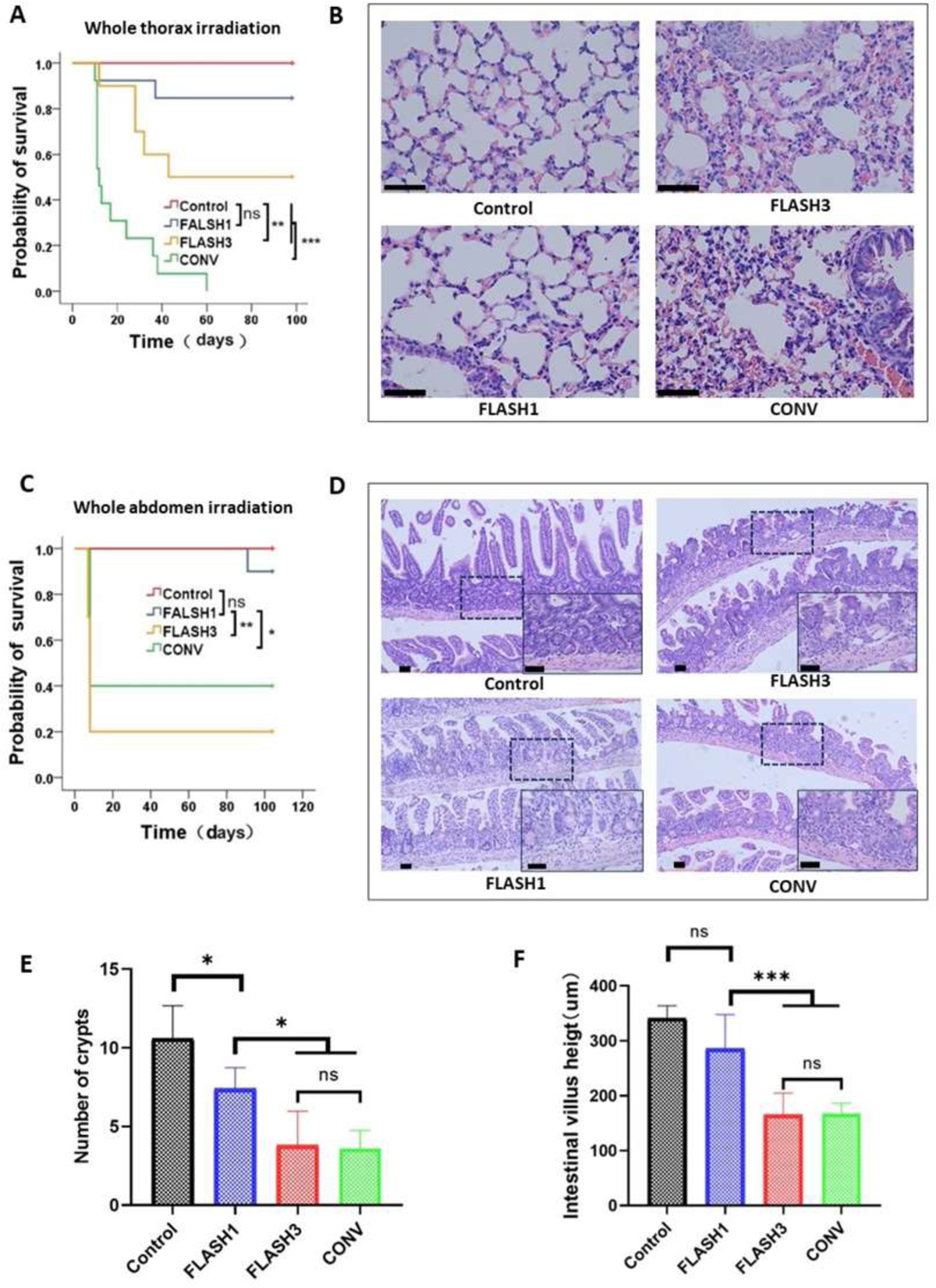
FLASH-RT minimizes lung and intestinal tissue damage. A: Survival curves of healthy C57BL/6 mice in the control (n=18), FLASH1 (30 Gy × 1F, 6 MeV, 340 Gy/s, n=18), FLASH3 (10 Gy × 3F, 6 MeV, 340 Gy/s, n=15), and CONV (30 Gy × 1F, 6 MeV, 0.07 Gy/s, n=18) groups after whole-thorax irradiation. B: H&E staining of the lung tissue under microscope(40x). C: Survival curves of healthy C57BL/6 mice in the control (n=15), FLASH1 (12 Gy ×1F, 6 MeV, 244 Gy/s, n=15), FLASH3 (4 Gy ×3F, 6 MeV, 244 Gy/s, n=15), and CONV (12 Gy × 1F, 6 MeV, 0.07 Gy/s, n=15) groups after whole-abdomen irradiation. D: H&E staining of the small intestine tissue under microscope(10x and 40x). E: Comparison among groups for the number of regenerating crypts within a single field of view (40x) under microscope. F: Comparison among groups for the height of small intestinal villi within a single field of view (40x) under microscope. *P<0.05, **P<0.01, ***P<0.001, and no significance (n.s.) were determined by a log-rank test (A, C) or one-way analysis of variance (E, F). Scale bars, 50 μm.

At 104 days post-whole-abdomen irradiation, the survival rates were 100%, 90%, 20%, and 40% in the control, FLASH1, FLASH3, and CONV groups, respectively (Fig. 2C). Survival rates were significantly higher in the FLASH1 group than those in the FLASH3 (P=0.001) and CONV (P=0.018) group, and similar between the FLASH3 and CONV (P=0.127) groups. In the intestinal tissues, H&E sections at 72 h post-irradiation revealed a significantly higher crypt number in the FLASH1 group than those in FLASH3 (7.40±1.34 vs. 3.80±2.17, P=0.019) and CONV (7.40±1.34 vs. 3.60±1.14, P=0.013) groups, while the crypt number was similar between the FLASH3 and CONV groups (3.80±2.17 vs. 3.60±1.14, P>0.999) (Fig. 2D, 2E). The intestinal villus height was significantly higher in the FLASH1 group than those in FLASH3 (285.41±62.27 vs. 165.41±39.37 µm, P<0.001) and CONV (285.41±62.27 vs. 167.03±19.24 µm, P<0.001) groups, with a similar intestinal villus height between the FLASH3 and CONV groups (165.41±39.37 vs. 167.03±19.24 µm, P>0.999) (Fig. 2F).

At 2 weeks post-skin irradiation, the incidence of ≥ grade 2 radiation dermatitis was 0%, 0%, 14.29%, and 85.71% in the control, FLASH1, FLASH3, and CONV groups, respectively (Fig. 3A, 3B). In the skin tissues, H&E sections at 8-week post-irradiation revealed a significantly higher epidermis thickness in the CONV group than those in control (232.99±113.97 vs. 12.95±2.35 um, P<0.0001), FLASH1 (232.99±113.97 vs. 31.03±9.54 um, P<0.0001), and FLASH3 (232.99±113.97vs. 60.66±40.21 um, P<0.0001) groups, with a similar epidermis thickness among the control, FLASH1, and FLASH3 groups (12.95±2.35 vs. 31.03±9.54 vs. 60.66±40.21 um, P>0.05) (Fig. 3C). Masson’s trichome staining at 8-weeks post-irradiation of skin tissue revealed a significantly higher percentage of collagen fiber deposition area in the CONV group than those in control (39.65±10.52 vs. 11.31±2.10, P<0.0001), FLASH1 (39.65±10.52 vs. 23.133±5.99, P<0.0030), and FLASH3 (39.65±10.52 vs. 25.36±11.31, P<0.0143) groups.

**Fig. 3:**
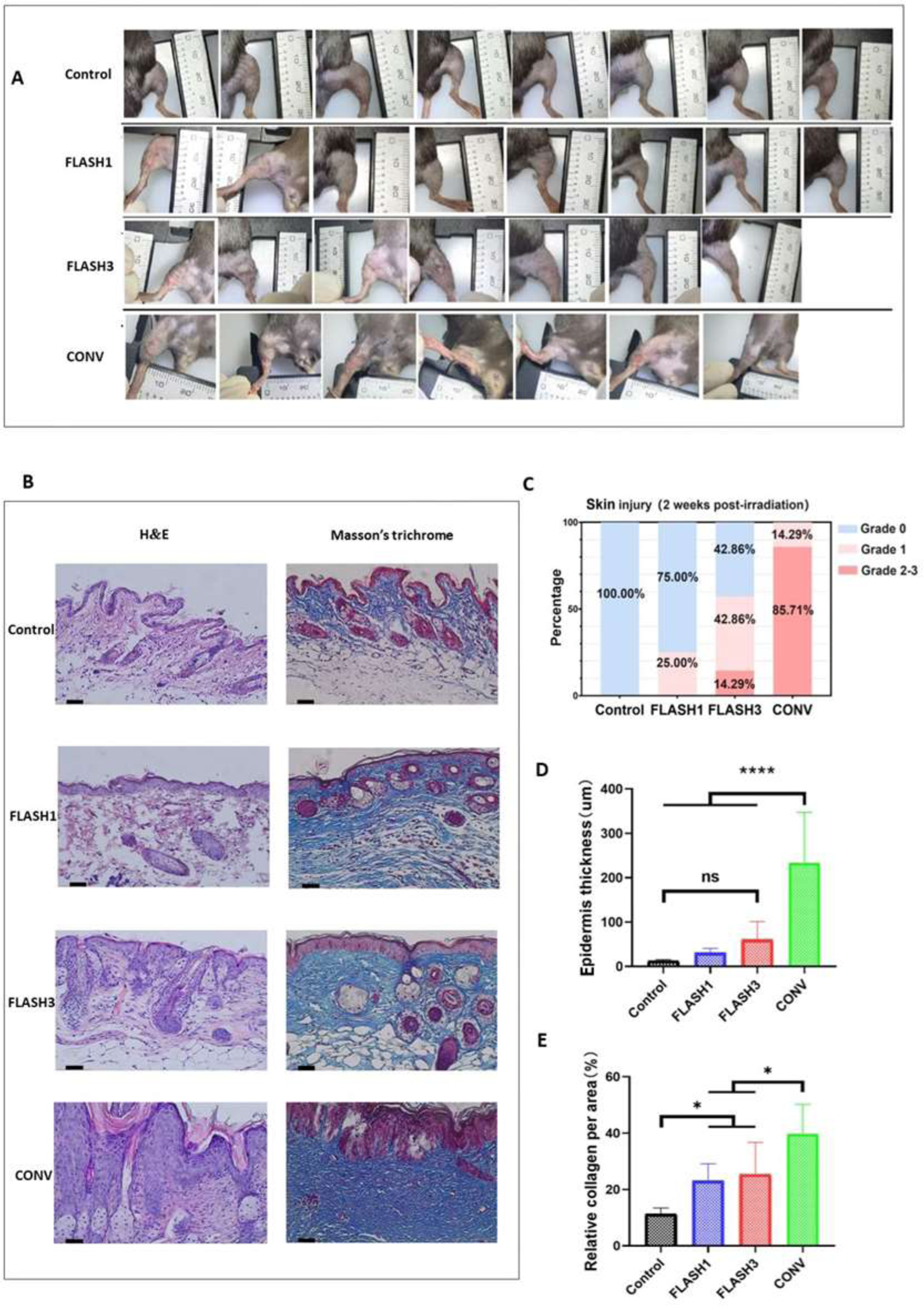
FLASH-RT minimizes skin tissue damage. A: Skin toxicities of healthy C57BL/6 mice in the control (n=8), FLASH1 (36 Gy × 1F, 6 MeV, 350 Gy/s, n=8), FLASH3 (12 Gy × 3F, 6 MeV, 350 Gy/s, n=7), and CONV (36 Gy × 1F, 6 MeV, 0.07 Gy/s, n=7) groups at 2 weeks post-irradiation. B: H&E and Masson’s trichrome staining of the skin tissue under microscope(10x). C: Incidence of skin toxicity (grades 0, 1, 2, and 3) among the groups. D: Comparison of epidermal thickness among the groups. E: Comparison of fibrosis among the groups. *P<0.05, ****P<0.0001, and no significance (n.s.) were determined by one-way analysis of variance (D, E). Scale bars, 50 μm.

Two tumor models (CT26 and LLC) were used to assess the anti-tumor effects of CHEXs. The tumor growth rates in the control group were higher than those in the FLASH1, FLASH3, and CONV groups (Fig. 4A, 4F).

**Fig. 4:**
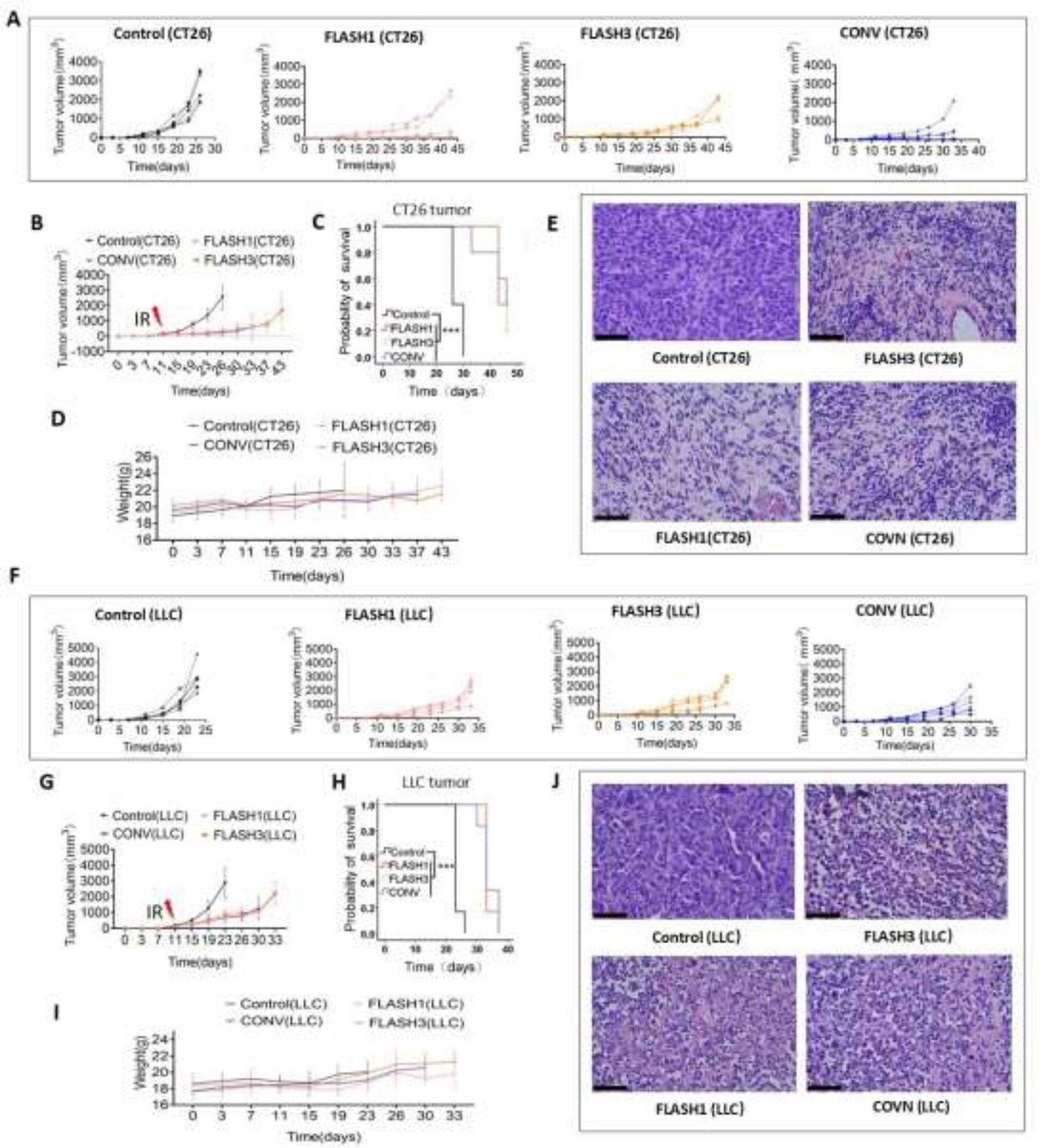
FLASH-RT retains anti-tumor efficacy. A, E, I: Tumor growth curves of the CT26, LLCmouse models. B, F, J: Mean CT26, LLC, and B16-F10 tumor growth curves among the groups. C, G: Image of mice with CT26 and LLC tumors taken on day 24 post-inoculation. K: Image of mice with B16-F10 tumors taken on day 19 post-inoculation. D, H, L: H&E staining of the CT26, LLC, and B16-F10 tumor tissues under microscope(40x). ****P<0.0001 and no significance (n.s.) were determined by one-way analysis of variance (D, E). Scale bars, 50 μm.

In the CT26 tumor model, tumor volumes were significantly larger in the control group than those in the FLASH1 (2564.73±793.89 vs. 343.24±305.91 mm^3^, P<0.0001), FLASH3 (2564.73±793.89 vs. 299.10± 60.74 mm^3^, P<0.0001), and CONV (2564.73±793.89 vs. 233.88±270.70 mm^3^, P<0.0001) groups at 15-day post-irradiation, with comparable tumor volumes among the FLASH1, FLASH3, and CONV groups (343.24±305.91 vs. 299.10±60.74 vs. 233.88±270.70 mm^3^, P>0.05) (Fig. 4B). The survival times were significantly longer in the FLASH1, FLASH3 and CONV groups than those in control group (P<0.001) (Fig. 4C). The weights of mice in four groups were similar (Fig. 4D). H&E staining revealed normal cellular morphology in the control group and tumor necrosis in the FLASH1, FLASH3, and CONV groups (Fig. 4E).

In the LLC tumor model, tumor volumes were significantly larger in the control group than those in the FLASH1 (2880.22±938.18 vs. 657.39±280.63 mm^3^, P<0.0001), FLASH3 (2880.22±938.18 vs. 857.05±347.50 mm^3^, P<0.0001), and CONV (2880.22±938.18 vs. 686.70±358.94 mm^3^, P<0.0001) groups at 12-day post-irradiation, with comparable tumor volumes among the FLASH1, FLASH3, and CONV groups (657.39±280.63 vs. 857.05±347.50 vs. 686.70±358.94 mm^3^, P>0.05) (Fig. 4G). The survival times were significantly longer in the FLASH1, FLASH3 and CONV groups than those in control group (P<0.001) (Fig. 4H). The weights of mice in four groups were similar (Fig. 4I). H&E staining revealed normal cellular morphology in the control group and tumor necrosis in the FLASH1, FLASH3, and CONV groups (Fig. 4J).

## Discussion

This study represents the first validation of the FLASH effect using the CHEXs device, which met the physical parameters for clinical application (equipment diameter approximately 3.1 meters, dose rate up to 81.01 Gy/s at 1 m SSD, and field diameter exceeding 10 cm) [13]. We also demonstrated that single gantry rotation two 30 s pauses irradiation can trigger the FLASH effect. Our findings validate the protective effect of CHEXs on normal tissues across multiple systems, including the lungs, intestines, skin, and bone marrow, and demonstrate that, compared with CONV-RT, FLASH-RT can significantly reduce whole-chest and whole-abdominal irradiation-related mortality rates in mice, the incidence of skin irradiation-induced grade 2 radiation dermatitis at 2-week post-irradiation, and bone marrow hematopoietic function damage. Upon comparing the anti-tumor efficacy of FLASH-RT and CONV-RT in mouse models of common malignant tumor types (lung cancer, colon cancer, and melanoma), we also confirmed that the high-energy X-rays produced by CHEXs exhibit the same anti-tumor efficacy as conventional dose-rate X-rays.

Current published [25,26] and ongoing (NCT05724875, NCT05524064, NCT04986696) clinical studies on FLASH-RT have mostly employed two-dimensional radiation therapy technology in patients with skin tumors and bone metastases for palliative pain relief purposes. However, in tumors with more complex shapes that implicate internal organs, three-dimensional (3D) conformal therapy techniques, such as FLASH-RT, may be necessary [27]. PHASER is a conceptual shared research platform designed to achieve highly conformal electronic FLASH-RT. The core concept behind PHASER is to utilize 16 coplanar heads to simultaneously output ultra-high-dose-rate electron beams, replacing single gantry rotation, to deliver 3D conformal FLASH-RT [28]. However, implementing multi-accelerator synchronous irradiation for high-energy X-ray sources remains challenging and expensive. Therefore, the present study explored whether single-gantry rotating two 30 s pauses irradiation can generate FLASH effects, thereby replacing the multi-accelerator synchronous irradiation mode. A three-stage segmentation mode simulating the three-field 3D conformal radiotherapy technique commonly used in clinical practice was constructed, with an interval of 30 s between each of the two radiotherapy sessions to replicate the time required for machine rotation. Our results confirmed that under the same total dose conditions, the FLASH effect can be observed in lung and skin tissues with two 30 s pauses irradiation (FLASH3). However, the protective effects of FLASH3 were diminished compared to FLASH, and no FLASH effect was observed in the intestinal tissue of the FLASH3 group. This may have been attributed to the single dose not meeting the threshold condition for triggering the FLASH effect (4 Gy), which has been reported to be approximately 10 Gy [24].

This study had several limitations. First, we used a single-gantry rotation mode instead of multi-gantry synchronous irradiation to achieve a 3D conformal dose distribution; however, because we used a mouse model, we set FLASH3 group included two 30 s pauses to simulate a three-field delivery where the gantry rotation is occurring within 30 s, rather than true single-gantry rotation 3D conformal radiotherapy. Nonetheless, in clinical radiotherapy, clinicians are most concerned with the normal tissue surrounding the tumor, which is often irradiated from multiple angles and has a relatively high dose distribution in 3D conformal radiotherapy.

Hence, the significance of our experimental results lie in the fact that the normal tissue in the high-dose area surrounding the tumor could also trigger the FLASH effect through two 30 s pauses irradiation.

## Conclusion

This study confirmed that the FLASH effect could be triggered using CHEXs FLASH radiotherapy, and demonstrated that three irradiation with single gantry rotation two 30 s pauses irradiation can trigger the FLASH effect, indicating the potential benefit of CHEXs 3D conformal radiotherapy. Our findings indicate that further clinical trials on CHEXs are warranted.

## Supplementary material

Parameters and dosimetry of the FLASH-RT and CONV-RT in whole-abdomen, skin and tumor irradiation.

## Acknowledgements

This work was supported by the Projects of National Natural Science Foundation (U2330122 and 12035012), Sichuan Natural Science Foundation General Program (2023NSFSC0710 and 2023NSFC3507), Postdoctoral Special Funding Research Project of Sichuan Provincial Department of Human Resources and Social Security (TB2023096).

