## Supplementary material for "FLASH radiotherapy using high-energy X-rays: validation of the FLASH effect triggered by a compact single high-energy X-ray source device"

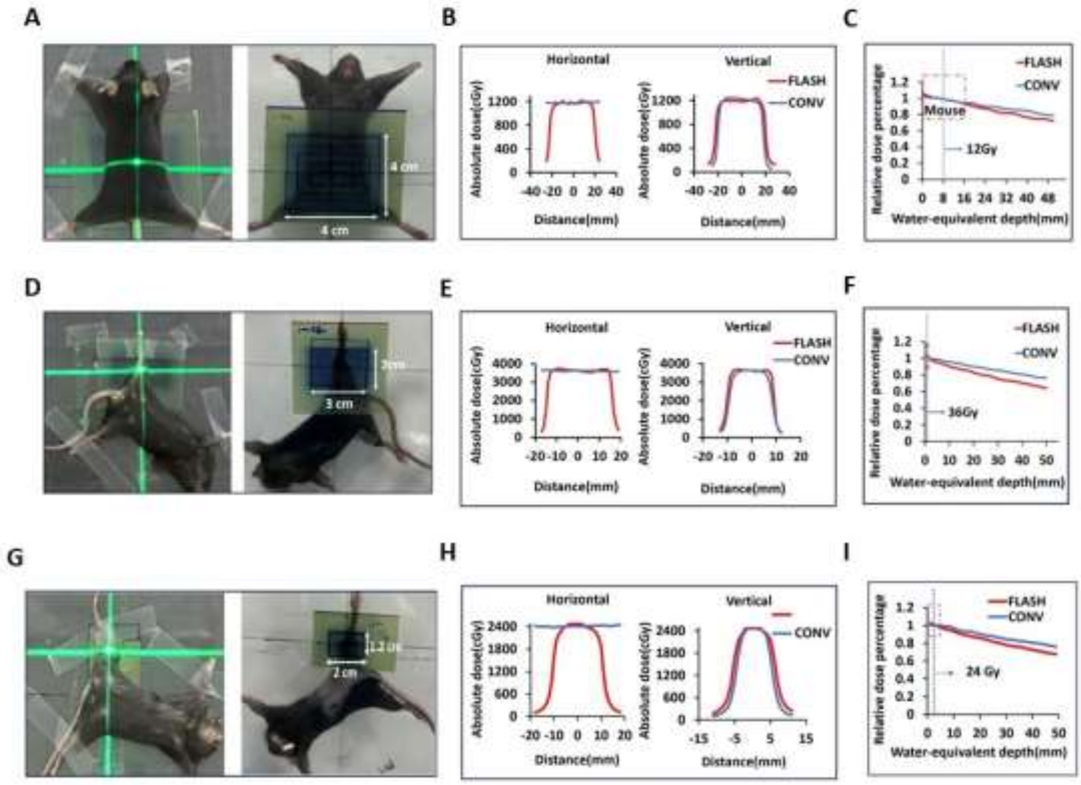


**Supplementary Fig. 1:** Parameters and dosimetry of FLASH-RT and CONV-RT in whole-abdomen, skin and tumor irradiation. A, D, G: FLASH-RT of whole-abdomen, skin and tumor irradiation, with 5-mm-thick polymethyl methacrylate (PMMA) plates used for build-up and mice fixing, and an EBT-XD film placed between the PMMA plate and the anterior surface of the irradiated mouse for dose evaluation. B, E, H: Horizontal and vertical dose profiles of FLASH-RT and CONV-RT in whole-abdomen, skin and tumor irradiation. C, F, I: The PDD FLASH-RT and CONV-RT curves in whole-abdomen, skin and tumor irradiation, with red dashed boxes representing the target areas.

Supplementary Table 1 The experimental parameters for whole-thorax, whole-abdomen, skin, whole-body and tumor irradiation.

| Experiment | mice | Filed size | | Number of mice | Beam energy | Dose delivered | Dose rate |
|---|---|---|---|---|---|---|---|
| Tumor irradiation | BAL b/c female mice, CT26 | 2 cm(lateral)×1.2 cm (craniocaudal) | Control | 8 | \ | \ | \ |
| | | | FLASH1 | 8 | 10 MeV | 16.5Gy*1F | 244 Gy/s |
| | | | FLASH3 | 8 | 10 MeV | 5.5 Gy*3F | 244 Gy/s |
| | | | CONV | 8 | 6 MeV | 16.5 Gy*1F | 0.07 Gy/s |
| | C57BL/6 female mice，LLC | 2 cm(lateral)×1.2 cm (craniocaudal) | Control | 9 | \ | \ | \ |
| | | | FLASH1 | 9 | 10 MeV | 18 Gy*1F | 244 Gy/s |
| | | | FLASH3 | 9 | 10 MeV | 6Gy*3F | 244 Gy/s |
| | | | CONV | 9 | 6 MeV | 18 Gy*1F | 0.07 Gy/s |
| Whole thorax irradiation | C57BL/6 female mice | 3 cm (lateral) ×2 cm (craniocaudal) | Control | 18 | \ | \ | \ |
| | | | FLASH1 | 18 | 10 MeV | 30 Gy*1F | 340 Gy/s |
| | | | FLASH3 | 15 | 10 MeV | 10 Gy*3F | 340 Gy/s |
| | | | CONV | 18 | 6 MeV | 30 Gy*1F | 0.1 Gy/s |
| Whole abdomen irradiation | C57BL/6 female mice | 4cm (lateral) ×4 cm (craniocaudal) | Control | 15 | \ | \ | \ |
| | | | FLASH1 | 15 | 10 MeV | 12 Gy*1F | 244 Gy/s |
| | | | FLASH3 | 15 | 10 MeV | 4 Gy*3F | 244 Gy/s |
| | | | CONV | 15 | 6 MeV | 12 Gy*1F | 0.07 Gy/s |
| Skin irradiation | C57BL/6 female mice | 3 cm (lateral) ×2 cm (craniocaudal) | Control | 8 | \ | \ | \ |
| | | | FLASH1 | 8 | 10 MeV | 36 Gy*1F | 350 Gy/s |
| | | | FLASH3 | 7 | 10 MeV | 12 Gy*3F | 350 Gy/s |
| | | | CONV | 7 | 6 MeV | 36 Gy*1F | 0.07 Gy/s |
